# GBS and a newly developed mRNA-GBS approach to link population genetic and transcriptome analyses reveal pattern differences between sites and treatments in red clover (*Trifolium pratense* L.)

**DOI:** 10.1101/2021.11.30.470612

**Authors:** B Gemeinholzer, O Rupp, A Becker, M. Strickert, C-M Müller

## Abstract

The important worldwide forage crop red clover (*Trifolium pratense* L.) is widely cultivated as cattle feed and for soil improvement. Wild populations and landraces have great natural diversity that could be used to improve cultivated red clover. However, to date, there is still insufficient knowledge about the natural genetic and phenotypic diversity of the species. Here, we developed a low-cost transcriptome analysis (mRNA-GBS) with reduced complexity and compared the results with population genetic (GBS) and previously published mRNA-Seq data, to assess whether analysis of intraspecific variation within and between populations and transcriptome responses is possible simultaneously. The mRNA-GBS approach was successful. SNP analyses from the mRNA-GBS approach revealed comparable patterns to the GBS results, but it was not possible to link transcriptome analyses with reduced complexity and sequencing depth to previously published greenhouse and field expression studies. The use of short sequences upstream of the poly(A) tail of mRNA to reduce complexity are promising approaches that combine population genetics and expression profiling to analyze many individuals with trait differences simultaneously and cost-effectively, even in non-model species. Our mRNA-GBS approach revealed too many additional short mRNA sequences, hampering sequence alignment depth and SNP recovery. Optimizations are being discussed. Nevertheless, our study design across different regions in Germany was also challenging as the use of differential expression analyses with reduced complexity, in which mRNA is fragmented at specific sites rather than randomly, is most likely counteracted under natural conditions by highly complex plant reactions at low sequencing depth.

## Introduction

*Trifolium pratense* (red clover) is an economically relevant crop in temperate agriculture, and also a major component of sustainable farming. *T. pratense* has a high protein content and serves as livestock fodder, promotes soil fertility and is an important component of crop rotation systems. Red clover is well known for its high biomass production and good re-growth capability after mowing (Kleen et al. 2011, Dewhurst 2013, Eriksen et al. 2014, Herbert et al. 2018). The species belongs to the Fabaceae (legumes) which encompasses several other agronomical important crops, like *Glycine max* (soy), *Medicago truncatula* (barrel clover), *Phaseolus vulgaris* (common bean), *Vigna unguiculata* (cowpea). *T. pratense* is diploid, which is important for high throughput molecular and functional analyses.

Agriculture is faced with the challenge of continuously optimizing crops in order to adapt them to changing climatic and cultivation conditions and to meet the steadily increasing demand for animal feed. In red clover, there is still high potential for breeding optimization, as in wild populations as well as in germplasm collections there exists a highly significant morphological and genetic variation (e.g. Dias et al. 2008, Kölliker et al 2003, Smith et al. 1985). The natural variability of the species, which is native to northwest Africa, throughout Europe, and much of Asia and has been introduced to North America, South America, Australia, and New Zealand, can be used in breeding programs to identify promising populations for improving agronomically important traits (e.g., plant size, growth habit, leaf area (Herbert et al. 2018), inflorescence size, number of inflorescences, flowering, disease susceptibility, and others (Isobe et al. 2009, Eriksen et al. 2014, Yates et al. 2014, deVega et al. 2015). This might especially be relevant in times of fast climatic and anthropogenic changes.

Today, rapidly evolving new NGS techniques, tools and analytical methods of genome and transcriptome sequencing, their statistical analysis and related informatics offer new opportunities to support agricultural breeding programs with genomic information. This allows for fostering knowledge in complex biological systems at various organizational levels (from individuals to populations, e.g. Wisecaver et al., 2017; Li et al., 2020), in different dimensions of time and space (Joly & Faure 2015 Gould et al., 2018; Mead et al., 2019; Marx et al. 2020) and under different treatments, greenhouse conditions, or in the field (Herbert et al., 2021). The development of the RNA-Seq method for quantitative next-generation sequencing of expressed genes has made expression studies for non-model species feasible. However, the method remains expensive and often requires a high number of replicates, so scalability is often not straightforward (Lohman et al., 2016). Genomic DNA fingerprinting (e.g., ddRadSeq; Hohenlohe et al. 2011) or genotyping by sequencing (GBS; Elshire et al. 2011)) is now widely used to perform association studies in many species, including those with complex genomes (Caballero et al. 2021), for revealing genetic diversity and population structure (Müller et al. 2019), for fingerprinting germplasm resources (Wang et al. 2021), or for the detection of candidate genes by fine mapping, especially for improving plant breeding strategies (e.g. Purugganan & Jackson 2021).

Here, we tested whether we can bridge the gap between genomic DNA fingerprinting and reduced complexity functional genomics in such a way that the natural diversity of a species can be studied quickly and inexpensively, so that the data can be linked relatively easily to functional analyses suitable for improving breeding programs. To achieve this, we developed an reduced complexity mRNA-GBS approach. We tested our mRNA-GBS approach on natural populations of red clover in three regions of the Biodiversity Exploratory sites in Germany, and evaluated how it correlates with genomic diversity of populations (analyzed with GBS) over a geographic range and to an earlier published gene expression profiling approach (mRNA-Seq, Herbert et al. 2021). Herbert et al. (2021) examined the expression patterns of red clover in relation to species-specific responses to mowing at one of the Biodiversity Exploratories and in the greenhouse. They identified candidate genes whose annotation suggests potential importance for phenotype changes in response to mowing. However, these analyses are currently only possible for a limited number of sites and individuals due to high costs and immense amounts of data (Gould et al. 2018; Marx et al. 2020). By combining fingerprinting with transcriptome profiling techniques across many samples, treatments, and locations, we test here whether it is possible to detect multiple genetic variants found across taxa and genomes in wild populations of red clover. Furthermore, we test whether this approach is able to simultaneously identify genomic population differences and candidate gene-signals potentially indicative for adaptive genetic variation. Our goal was to assess whether mRNA-GBS provides results that are equitable and relatable to GBS and RNA-Seq, are biologically informative, and are more cost-effective due to the shallow sequencing depth.

## Materials & Methods

### Study site and sampling

Sampling of plant material for mRNA-GBS and GBS was performed on the premises of the long-term open research platform Biodiversity Exploratory in June 2017 on the three Biodiversity Exploratories “Schorfheide-Chorin (S)” in the State of Brandenburg, “Hainich-Dün (H)” in Thuringia, and “Swabian Alb (A)” in the State of Baden-Württemberg, Germany (Fischer et al. 2010) at six field sites each (Table 1, Fig. 1). One population (AEG9) deviated so much from the other populations in its values and patterns that it was excluded from further analyses in the mRNA-GBS as well as in the GBS analysis. The experimental plots were managed as normal agricultural land colonized with native, established red clover populations. The not-mown pastures and meadows were neither grazed nor mown in the year of sampling (Herbert et al. 2021). Collection permits from farmers and local authorities were obtained centrally by the Biodiversity Exploratory research platform. At least seven individuals per site (126 in total) were quick-frozen in liquid nitrogen in the field and stored at - 80°C until further processing.

**Fig. 1.**
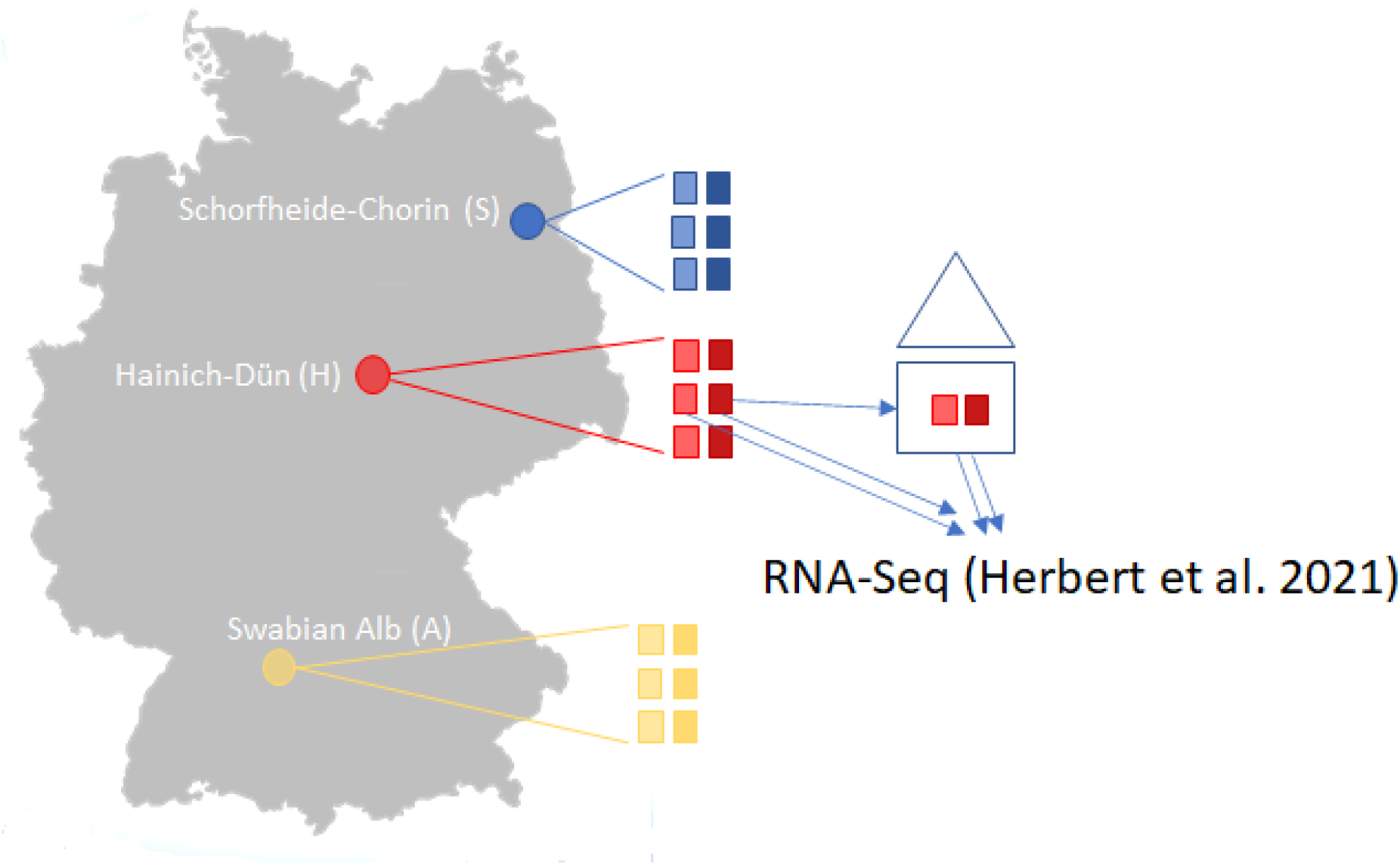
Study sites in Germany of the three Biodiversity Exploratory sites (S: Schorfheide-Chorin; H: Hainich-Dün; A: Swabian Alb) with 6 sampled populations per site (three mown (transparent colors) and three unmown (rich colors)) for the mRNA-GBS analysis and the GBS analysis and results were compared to the RNA-Seq-study of Herbert et al. (2020) where samples derived from the Hainich-Dün site directly and were cultivated in a greenhouse experiment.

**Table 1.** Study sites

| Region | Patch ID/Plot ID | Treatment | Date of mowing | Date of sampling | Sampling days after mowing | lat | long | mRNA-GBS/GBS |
| --- | --- | --- | --- | --- | --- | --- | --- | --- |
| Schorfheide-Chorin (S) | SEG30/S690 | mown | 17.06.2017 | 11.07.2017 | 25 | 53,1481413 | 13,8306805 | 7/5 |
|  | SEG31/S698 | mown | 17.06.2017 | 11.07.2017 | 25 | 53,1490865 | 13,8354379 | 7/5 |
|  | SEG32/S701 | mown | 17.06.2017 | 11.07.2017 | 25 | 53,1519293 | 13,8320874 | 7/5 |
|  | SEG z31-32 | not mown | - | 11.07.2017 | - | 53,150246 | 13,834948 | 7/5 |
|  | SEG HG | not mown | - | 11.07.2017 | - | 53,147784 | 13,834846 | 7/5 |
|  | SEG z30-31 | not mown | - | 11.07.2017 | - | 53,14827 | 13,833452 | 7/5 |
| Hainich-Dün (H) | HEG8/H14707 | not mown | - | 16.06.2017 | - | 51,2712575 | 10,4179446 | 7/5 |
|  | HEG13/H4651 | mown | 23.05-01.06.2017 | 16.06.2017 | 16-23 | 51,2596811 | 10,3799461 | 7/5 |
|  | HEG15/H16781 | mown | 15.06.2017 | 11.07.2017 | 28 | 51,068013 | 10,4862323 | 7/5 |
|  | HEG17/H14529 | not mown | - | 16.06.2017 | - | 51,0705045 | 10,4704561 | 7/5 |
|  | HEG50/H15457 | not mown | - | 16.06.2017 | - | 51,2765094 | 10,4207739 | 7/5 |
|  | HEG50/H15457 | mown | 13.06.2017 | 11.07.2017 | 30 | 51,2765094 | 10,4207739 | 7/5 |
| Swabian Alb (A) | AEG2/A39275 | mown | 26.05.2017 | 14.06.2017 | 20 | 48,3768573 | 9,47278412 | 7/5 |
|  | AEG14/A46088 | not mown | - | 14.06.2017 | - | 48,37576 | 9,51866928 | 7/5 |
|  | AEG15/A35767 | mown | 25.05.2017 | 14.06.2017 | 21 | 48,4878818 | 9,44865392 | 7/5 |
|  | AEG24/A42306 | mown | 25.05.2017 | 14.06.2017 | 21 | 48,3964649 | 9,49349225 | 7/5 |
|  | AEG31/A37367 | not mown | - | 14.06.2017 | - | 48,4587449 | 9,46002446 | 7/5 |

Region, Patch ID/Plot ID according to Fischer et al. 2010, agricultural treatment with dates and sampling dates, geographic locations with latitudes (lat) and longitudes (long), sampled individuals (n) for the mRNA-GBS and the GBS analysis.

### Molecular techniques

Briefly, our mRNA-GBS library construction method involves 8 laboratory steps: (i) isolate total RNA, (ii) remove genomic DNA with DNase, (iii) convert mRNA into cDNA by using a reverse transcription kit (cDNA) using a BceA restriction sites containing PolyA primer with an anchor, (iv) digestion with BceA and MseI restriction enzymes, (v) NGS primer ligation with BceA adapter and index and MseI adapter, (vi) pooling, purification and PCR amplification, (vii) size selection, (viii) Illumina Next Seq 500 Vs sequencing (Fig. 2).

**Fig. 2.**
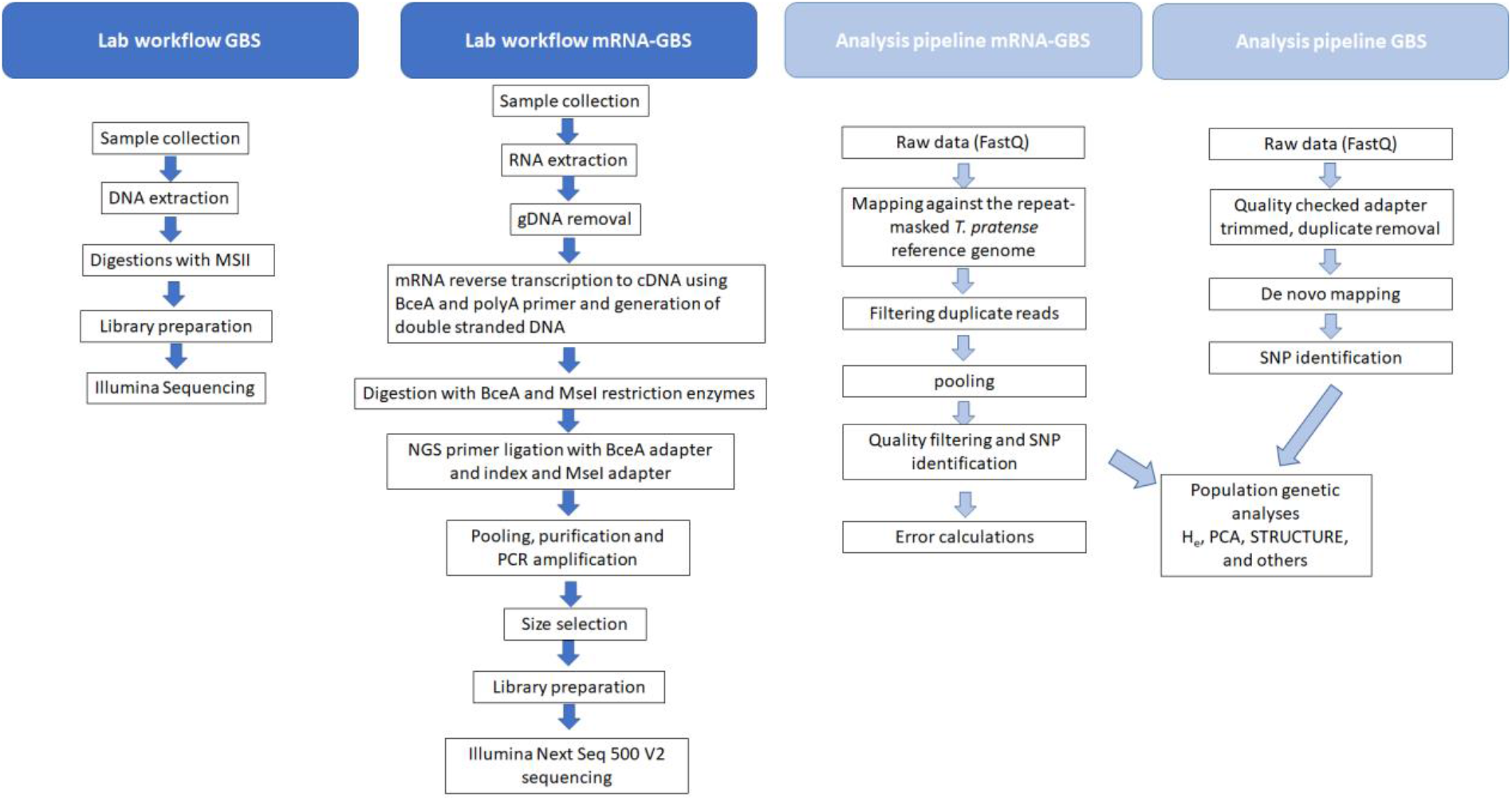
Laboratory and data analysis workflow

For the mRNA-GBS analysis seven individuals per site were examined. For RNA extraction we used the NucleoSpin^®^ RNA Plant kit (Macherey-Nagel, Germany) according to the manufacturer’s instructions. For the mRNA-GBS development the Maxima H Minus Double-Stranded cDNA Synthesis Kit (Thermo Scientific™, Germany) was used for double stranded cDNA-Synthesis, however, with a specially designed PolyT priming site, suitable to be cleaved by the BceAI restriction enzyme (gcBceAI-PolyA-TVN-Primer: 5’-CCGGCGCGACGGCTTTTTTTTTTTTTTTTVN-3’) following the user manual. Purification took place with the NucleoSpin^®^ Gel and PCR Clean-up kit (Macherey-Nagel, Germany). Restriction was carried out, by digesting 200 ng double stranded cDNA with BceAI (2 U/μl) and MseI (10 U/μl) by 37°C in NEB 3.1 buffer (16μl cDNA/H_2_O (200ng cDNA), 2μl buffer, 1μl BceAI, 0.25μl MseI and 0.75μl H_2_O.60 min incubation at 37°C and 20 min inactivation at 65°C). After preparing the samples 30ng/μl of the digested material were transferred to LGC Genomics GmbH (Germany) for library preparation, pooling and sequencing (150 bp paired-end reads on an Illumina Next Seq 500 V2, Fig. 2).

For GBS analysis, DNA was extracted from five samples per site (Table 1). We used the Invisorb^®^ Spin Plant Mini Kit from Stratec Molecular (Germany) according to the instructions for use. DNA quantity and quality were analysed using a NanoPhotometer™ (Implen GmBH, München, Germany). We sent 300ng of DNA in 20μl to LGC Genomics GmbH (Germany) where genomic DNA were digested with 1 Unit MslI (NEB) in 1 times NEB4 buffer in 30 μl volume for 1 h at 37 °C. The restriction enzyme was heat inactivated by incubation at 80 °C for 20 min. The indexed Illumina libraries were prepared by using the Encore Rapid Multiplex System (Nugen): 15 μl were transferred to a new 96 well PCR plate, mixed on ice first with 3 μl of one of the 192 L2 Ligation Adaptors and then with a 12 μl Mastermix (a combination of 4.6 μl D1 water/ 6 μl L1 Ligation Buffer Mix/ 1.5 μl L3 Ligation Enzyme Mix). Ligation reactions were incubated at 25 °C for 15 min and heat inactivated at 65 °C for 10 min. A 20 μl Final Repair Master Mix was added to each tube and the reaction was incubated at 72 °C for 3 min. For purification, the reactions were diluted with 50 μl TE 10/50 (10mM Tris/HCl, 50mM EDTA, pH: 8.0) and mixed with 80 μl Agencourt XP beads, incubated for 10 min at RT and placed for 5 min on a magnet to collect the beads. The supernatant was discarded and the beads were washed two times with 200 μl 80% Ethanol. Beads were air dried for 10 min and libraries were eluted in 20 μl Tris Buffer (5mM Tris/HCl pH:9) prior to sequencing on an Illumina NextSeq 500 V2, resulting in 150 bp paired-end reads.

### Bioinformatics and Genotyping

#### mRNA-GBS data SNP calling

The Illumina reads were mapped to the repeat-masked *T. pratense* reference genome (version GCA_900079335.1, ENSEMBL release 50) using the STAR short read mapper (Dobin and Gingeras, 2015). Duplicate reads were filtered using the Picard Toolkit (Broad Institute, 2019) MarkDuplicates algorithm (version 2.26.1). The samples of the same field site were pooled to get a higher resolution. Alleles were counted using bam-readcount (The McDonnell Genome Institute, 2021) with a minimum base quality of 20. Only loci with at least ten reads in each pool were considered and alleles were called only when supported by at least three reads. Error rates with TPM normalized read-counts were calculated using the following pipeline: http://rseqc.sourceforge.net/#rpkm-saturation-py

#### GBS data analysis

after base calling and demultiplexing the quality of the sequenced reads were quality checked. SNP calling and genotyping was conducted with Freebayes (Garrison and Marth 2021). We used adapter clipped data for further calculations in Stacks 1.48 (Catchen et al. 2011; Catchen et al. 2013). UStacks and denovo_map were applied for analyses without a reference genome. The following (default) parameters for the formation of stacks and loci were used: minimum depth of coverage to create a stack –m = 3, maximum of distance allowed between stacks –M = 2, distance allowed between catalog loci –n = 0, (maximum distance allowed to align secondary reads –N = 4, maximum number of stacks allowed per de novo locus: 3) and –t to remove or break up highly repetitive RAD-Tags in UStacks. Next we ran CStacks (to build the catalog) and SStacks (match the samples against the catalog) pipelines without modifications. We applied the correction module rxstacks, filtering by locus log likelihood with the following options: –t 40 --conf_lim 0.25 --prune_haplo --model_type bounded --bound_high 0.1 --lnl_lim −8.0 --lnl_dist –verbose. Finally, we ran the population program in Stacks with following parameters for: –r = 0.75. PGDSPIDER v.2.1.0.0 (Lischer and Excoffier 2012) was used to convert Stacks output files for further analyses.

Genetic diversity was estimated as percentage of polymorphic loci (PL) and as Nei’s gene diversity (H_e_; Nei (1973)) using ARLEQUIN v.3.5.1. (Excoffier and Lischer 2010) and the package “diveRsity” (Keenan et al. 2013) by using R 3.5.1 (R Core Team 2013). To visualize the data STRUCTURE (Pritchard et al. 2000) was used, which shows the membership probabilities. For automation and parallelization of STRUCTURE (Pritchard et al. 2000) analysis we used the program StrAuto (Chhatre and Emerson 2017). Genetic clusters were detected by applying the admixture model, with 1000 Markov Chain Monte Carlo (MCMC) replicates, with a burn-in period of 1000 and ten repeats per run for each chosen cluster number (i.e. K = 1 – 20), Ploidy = 2. For all other settings, default options were used. To identify the most likely K modal distribution, delta K (Evanno et al. 2005) was determined by using STRUCTURE HARVESTER (Earl and von Holdt 2012) wich is also integrated in StrAuto (Chhatre and Emerson 2017). To verify the most probable cluster membership coefficient among the ten runs of STRUCTURE and STRUCTURE HARVESTER we used CLUMPP v.1.1.2 (Jakobsson and Rosenberg 2007). Corresponding graphs were constructed with DISTRUCT (Rosenberg 2004). By using R 3.5.1 (R Core Team 2013) and the R package ‘adegenet’ v.1.4-2 (Jombart 2008) a Principal Component Analysis (PCA) was calculated. With the R package ‘adegenet’ v.1.4-2 (Jombart 2008) and ‘ape’ (Paradis et al. 2004) the dendrograms were calculated, euclidian distance was used. Genetic variation among groups of populations (F_CT_), among populations within groups (F_SC_) and within populations (F_ST_) were partitioned with hierarchical analyses of molecular variance (AMOVA) by using ARLEQUIN v.3.5.1.2 (Excoffier and Lischer 2010) with an allowed missing data level at 5 %. Additionally, pairwise F_ST_ values were estimated among populations, with significance levels of 0.05 and 100 permutations.

## Results

### Sampling and genotyping

The mRNA-GBS sequencing yielded a total of 183.747.290 reads for the 126 investigated samples, with 42 individuals per region (S, H, A; Table 2). Retrieved read numbers varied strongly between individuals with an average of 1.1 million raw reads per sample (range: 7.106.704 – 31.481). After applying different filtering steps, 91.870.548 adapter clipped read pairs were retrieved. To analyze error rates, we calculated TPM-normalized read counts for each sample (Fig. 3) by testing our mRNA-GBS library against the RNA-Seq library of Herbert et al. (2021). Since TPM normalizes to sequencing depth, the value should be stable with respect to the actual read count if the sequencing depth was appropriate. When we reduced our samples from 90% to 60% sequencing depth (Fig. 6), the changes in error rate indicated that our sequencing depth was insufficient to perform gene expression studies and to be matched against the *T. pratense* transcriptome (Herbert et al., 2021) for subsequent analysis, whereas the error rate in Herbert et al. (2021) was stable and in line with expectations.

**Fig. 3.** Error rates for the TPM normalized read counts for the samples of the mRNA-GBS analysis, depicted in light green (S), bluish green (H) and purple (A) and the RNA-Seq data of Herbert et al. (2020) in red, calculated with 90% coverage (upper left), 80% coverage (upper right), 70% (lower left), 60% (lower right) an revealing strong differences in the error rate detection in the mRNA-GBS samples, when coverage is reduced, with little differences in the RNA-Seq data, which is stable and thus usable for gene expression analysis.

To identify SNPs for population genetic studies, the sequencing depth for SNP analysis of individual samples was also too shallow. Therefore, individuals within sites of similar treatments (mown/not mown) were combined in bulk samples to obtain a site-specific pattern. In this way, a total of 15.111 SNPs were obtained for subsequent analysis.

**Table 2:** Number of raw reads retained in mRNA-GBS and GBS analysis after each filtering step for *Trifolium pratense* samples from the three Biodiversity Exploratory sites in Germany (S: Schorfheide-Chorin; H: Hainich-Dün; A: Swabian Alb).

|  | mRNA-GBS |  |  |  | GBS |  |  |  |
| --- | --- | --- | --- | --- | --- | --- | --- | --- |
| Sample name | Raw total reads | Raw read pairs | Adapter clipped reads | Adapter clipped read pairs | Raw total reads | Raw read pairs | Restriction enzyme filtered reads | Restriction enzyme filtered read pairs |
| Sum total | 183.747.290 | 91.873.645 | 183.741.096 | 91.870.548 | 296.844.208 | 148.422.104 | 290.780.210 | 145.390.105 |
| Sum S | 92.964.904 | 46.482.452 | 92.961.812 | 46.480.906 | 109.954.908 | 54.977.454 | 107.658.878 | 53.829.439 |
| Sum H | 64.219.776 | 32.109.888 | 64.217.018 | 32.108.509 | 103.920.304 | 51.960.152 | 101.930.726 | 50.965.363 |
| Sum A | 26.562.610 | 13.281.305 | 26.562.266 | 13.281.133 | 82.968.996 | 41.484.498 | 81.190.606 | 40.595.303 |

The GBS sequencing yielded a total of 296.844.208 raw reads (range: 2.212.232 – 777.242) for the 90 investigated samples from the three regions each (Table 1, Figure 1), on average 3.6 million reads per sample. After applying different filtering steps, 56.395 SNPs were obtained for subsequent analyses, which is an 3.7 times higher coverage than received via the mRNA-GBS analysis.

The mRNA-GBS analysis revealed a comparatively high mean genetic diversity of the investigated red clover bulk samples of ØH_e_ = 0.76, ranging from H_e_ = 0.72 (S) to H_e_ = 0.82 (A, Table 3), if the regions are to be considered. The genetic diversity is higher, if sites with treatments (mown/not mown) are to be considered ØH_e_ = 0.82, ranging from H_e_ = 0.79 (S mown) to H_e_ = 0.86 (A not mown, Table 3). Because the analysis included multiple combined individuals from three populations per site and only two sites per region, the population comparison was too low to calculate genetic diversity among regions. The GBS analysis revealed a significantly lower mean genetic diversity of the investigated red clover populations of ØH_e_ = 0.060, ranging from H_e_ = 0.049 (AEG31, AEG24) to H_e_ = 0.060 (HEG8, HEG13, Table 3). The region specific mean genetic diversity is lowest in A (ØH_e_ = 0.050), intermediate in S (ØH_e_ = 0.055) and highest in H (ØH_e_ = 0.058). According to the ANOVA, genetic diversity among the three regions differed significantly (ANOVA *F* = 9.255 *P* = 0.009). Tukey test showed a significant difference between A – H (*P* = 0.007) but not between H - S (*P* = 0.470) and A - S (*P* = 0.139). The ANOVA with polymorphic loci only revealed no differences between A, H and S (F = 2.731, *P* = 0.0997). The AMOVA revealed moderate genetic differentiation among regions (F_CT_ = 0.05) and within populations (F_ST_ = 0.07) which are highly significant. However, for among populations within regions the genetic differentiation is negligible (F_SC_ = 0.02, Table 3). Thus, differentiation within populations were greater than among regions. Pairwise population FST estimates for the entire study area indicates low genetic differentiation among populations (0.00 - 0.013, Figure X). Pairwise population differentiation within regions is low to negligible for all regions (Ø A FST = 0.01, Ø S FST = 0.022, Ø H FST = 0.016).

**Table 3:** Population genetic statistics

| Regions | Patch ID | Treatment | mRNA-GBS |  |  |  |  |  | GBS |  |  |
| --- | --- | --- | --- | --- | --- | --- | --- | --- | --- | --- | --- |
|  |  |  | H<sub>e</sub>-value | PL | PL% | H<sub>e</sub>-value | PL | PL% | H<sub>e</sub>-value | PL | PL% |
| Schorfheide-Chorin<br>(S) | SEG30 | mown |  |  |  |  |  |  | 0.057 | 7896 | 19.25 |
|  | SEG31 | mown | 0.79 | 10934 | NaN |  |  |  | 0.056 | 6717 | 18.57 |
|  | SEG32 | mown |  |  |  | 0.72 | 15615 | NaN | 0.059 | 7177 | 20.39 |
|  | SEGHG | not mown |  |  |  |  |  |  | 0.053 | 5475 | 16.99 |
|  | SEGz1 | not mown | 0.80 | 10625 | NaN |  |  |  | 0.054 | 5810 | 17.37 |
|  | SEGz2 | not mown |  |  |  |  |  |  | 0.053 | 5413 | 17.91 |
| Hainich-Dün<br>(H) | HEG13 | mown |  |  |  |  |  |  | 0.060 | 8565 | 20.34 |
|  | HEG15 | mown | 0.81 | 9050 | NaN |  |  |  | 0.059 | 8287 | 20.24 |
|  | HEG50g | mown |  |  |  | 0.75 | 13137 | NaN | 0.058 | 7617 | 20.08 |
|  | HEG17 | not mown |  |  |  |  |  |  | 0.051 | 5529 | 17.17 |
|  | HEG50 | not mown | 0.79 | 10950 | NaN |  |  |  | 0.058 | 7400 | 20.15 |
|  | HEG8 | not mown |  |  |  |  |  |  | 0.060 | 8317 | 20.79 |
| Swabian Alb<br>(A) | AEG2 | mown |  |  |  |  |  |  | 0.054 | 6739 | 18.58 |
|  | AEG15 | mown | 0.84 | 7215 | NaN |  |  |  | 0.050 | 7140 | 17.17 |
|  | AEG24 | mown |  |  |  | 0.82 | 8776 | NaN | 0.049 | 6781 | 16.77 |
|  | AEG14 | not mown |  |  |  |  |  |  | 0.050 | 6721 | 17.06 |
|  |  |  | 0.86 | 5711 | NaN |  |  |  |  |  |  |
|  | AEG31 | not mown |  |  |  |  |  |  | 0.049 | 6744 | 16.98 |

Regions (S: Schorfheide-Chorin; H: Hainich-Dün; A: Swabian Alb), Collection sites (Patch ID), Nei's gene diversity (H<sub>e</sub>), polymorphic loci (PL), total no. of SNP counts, percentage of polymorphic loci (PL%).

STRUCTURE analyses based on the BIC and Bayesian clustering approaches revealed two genetic clusters, the proportional cluster membership of each being almost region-specific in the GBS analysis (Fig. 4A). The mRNA-GBS approach resulted in similar trends that were less prominent (Fig. 4B). This is also confirmed by the PCA (Fig. 5), which shows the respective site specificity of the centroids of all individuals (GBS) or bulk samples (mRNA-GBS) belonging to one sampling region, however, with much greater genetic similarity between individuals from S and H and the greater distance from A in the GBS analysis and more overlap in the mRNA-GBS data. This overlap is partly due to mowing treatment: the mown populations in the mRNA-GBS analysis showed a stronger pattern of site specificity, while the mRNA-GBS pattern of the unmown individuals was highly divergent. The GBS Neighbor Joining tree (Fig. 6A) reflects the patterns of the AMOVA, PCA, and STRUCTURE analyses, with individuals from A distinctly different from those from H and S, with some minor overlap between H and S among the individuals considered. The mRNA-GBS tree (Fig. 6B) also reflects the separate positions of the populations in A, but shows more mixing between H and S. The not mown populations A (AEG31, AEG14, Fig. 6B), and two out of three of the not mown populations in S (SEGHG, SEGz1) are also clustered, but lack a clear pattern as several other not mown populations appear scattered in the tree (SEGz2, HEG17, HEG8, HEG50).

**Fig. 4.**
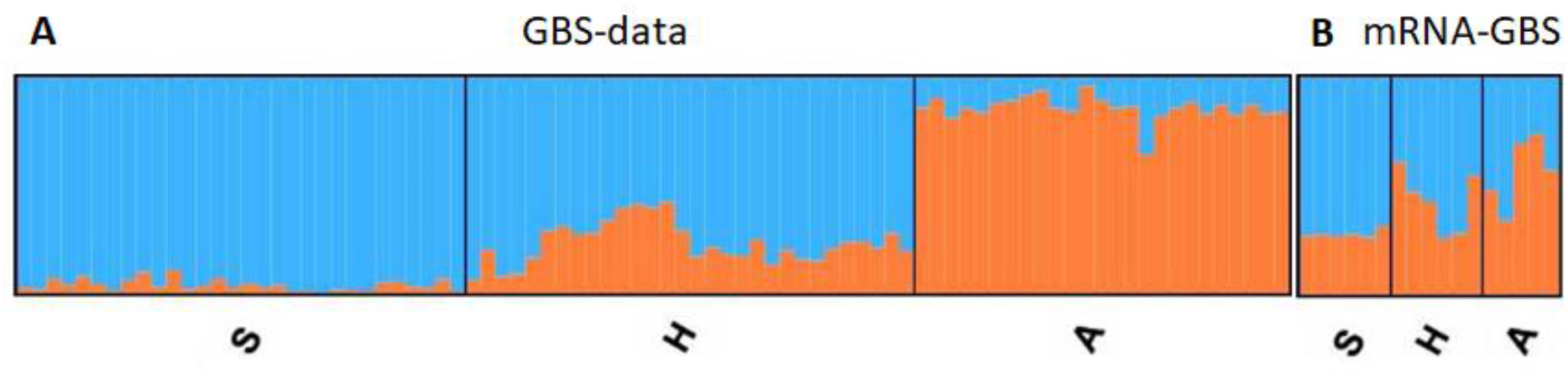
Population genetic structure of the investigated red clover individuals (GBS) or site specific bulk samples (mRNA-GBS) across the different Biodiversity Exploratories (S: Schorfheide-Chorin; H: Hainich-Dün; A: Swabian Alb) as revealed by the STRUCTURE analyses and ΔK (Evano et al. 2005). **A:** for the GBS data where each column represents individuals within one region; **B:** for mRNA-GBS data, where each column represents the bulk samples within one population.

**Fig. 5.**
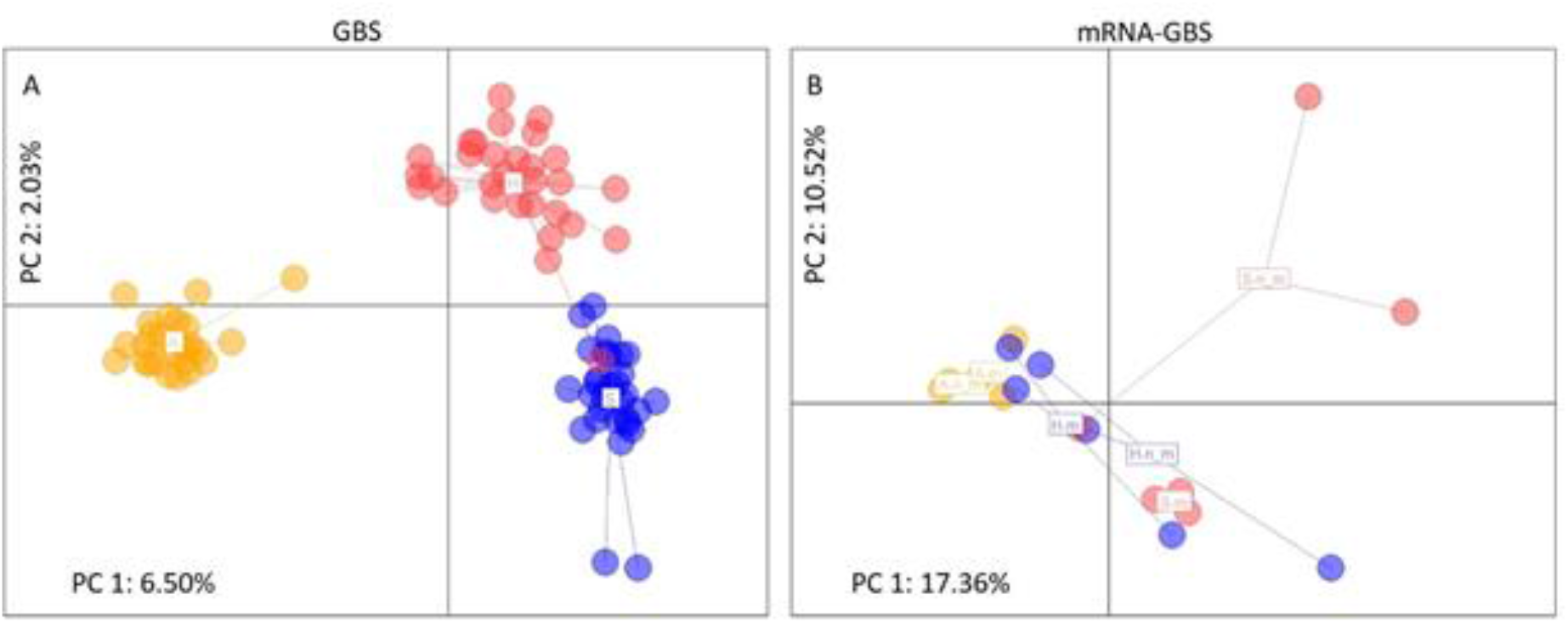
Principal Component Analysis (PCA) of genetic distances between individuals (GBS) or site specific bulk samples (mRNA-GBS) of *Trifolium pratense* across the different Biodiversity Exploratories (S: Schorfheide-Chorin; H: Hainich-Dün; A: Swabian Alb). Colored label positions represent the centroids of all individuals belonging to one sampling region for **A:** the GBS analysis, depicting colour coded individuals within each region, where the third axis is representing 1.85% of genetic variation (Σ 10.38%) and **B:** the mRNA-GBS analysis, depicting colour coded populations of bulk samples within each region (S, H, A where n_m is not mown, m is mown). The third axis is representing 8.72 % genetic variation (Σ 36.60%).

**Fig. 6.**
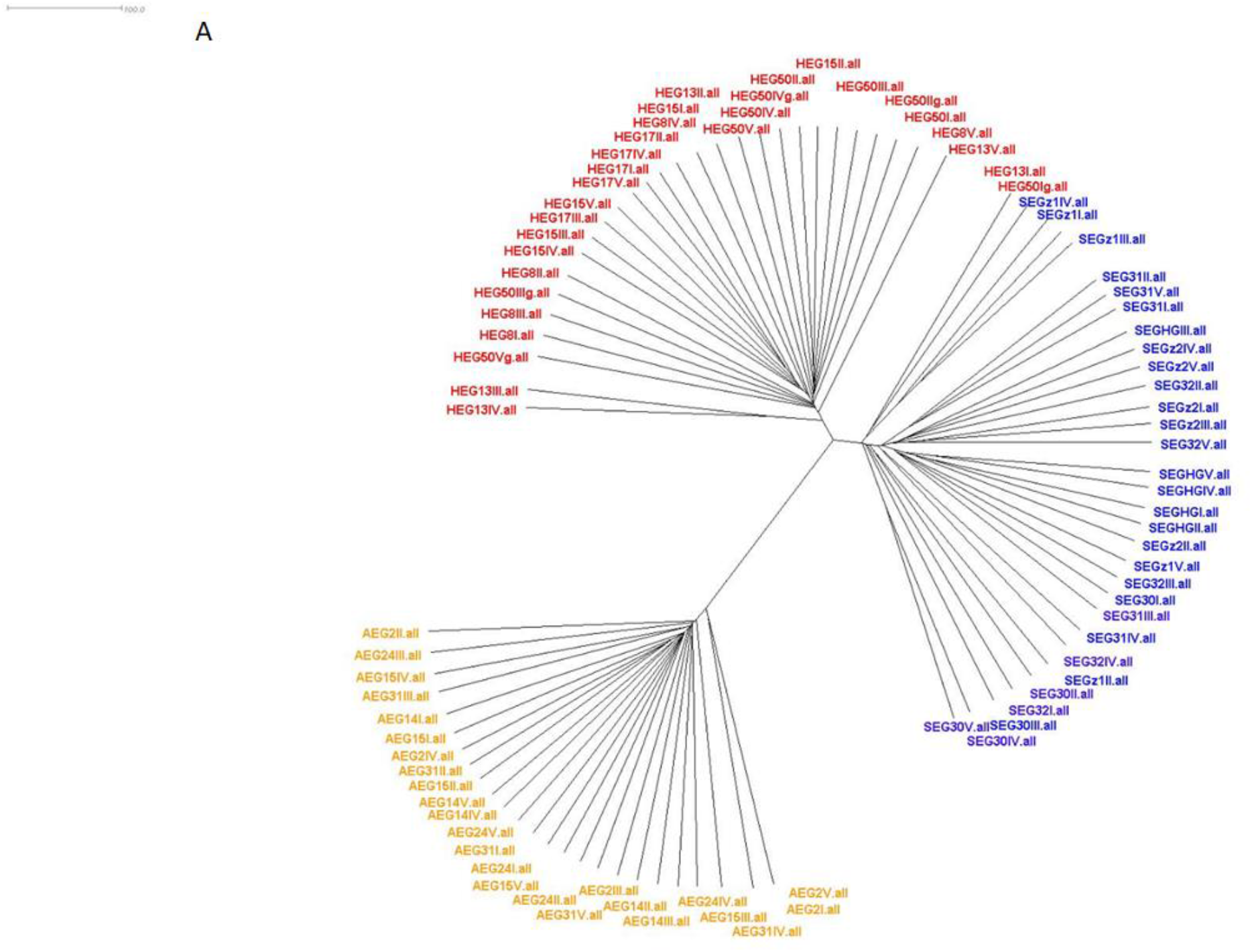
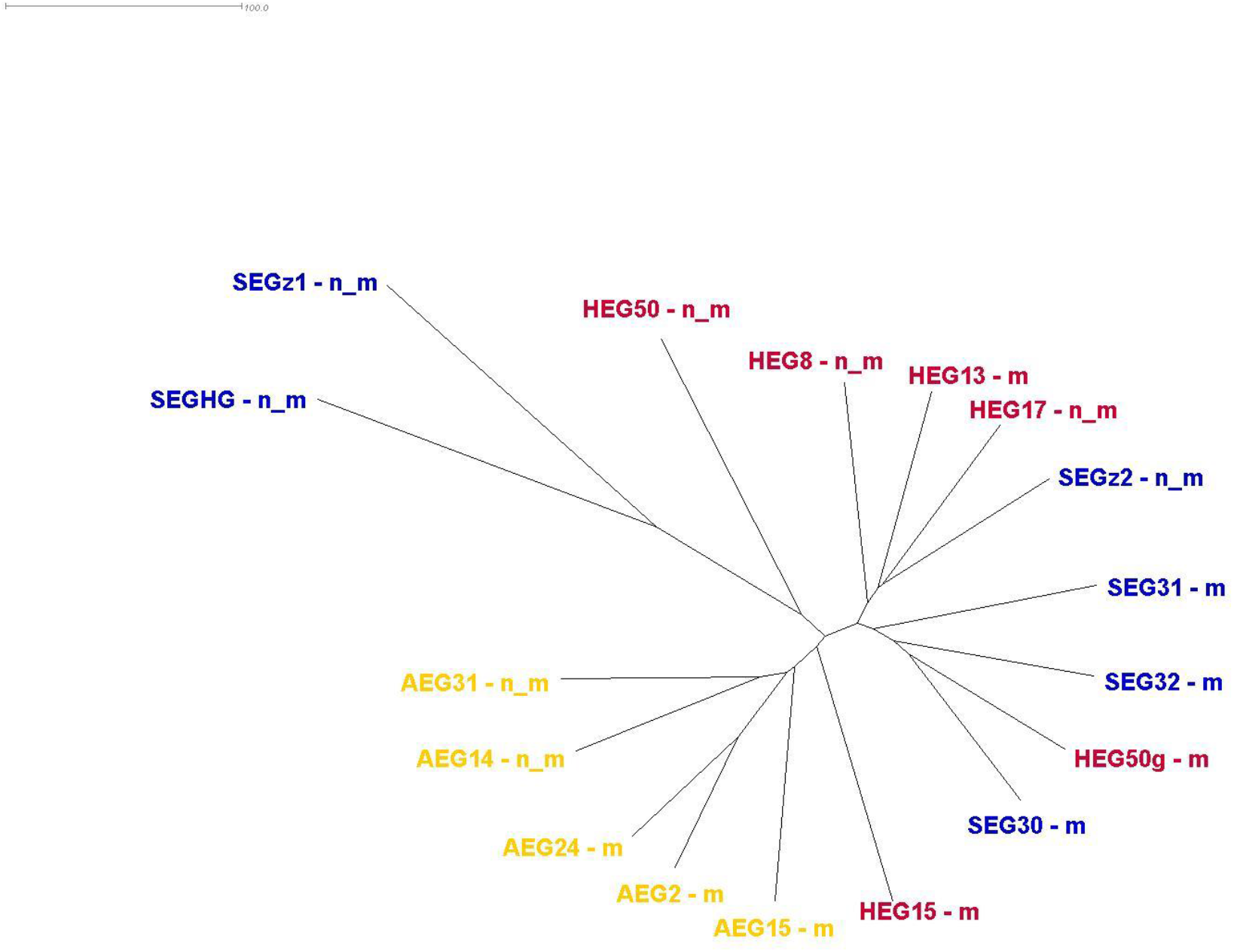
Neighbor Joining tree for the individuals and populations of *Trifolium pratense* across the different Biodiversity Exploratories (yellow: Schorfheide-Chorin; red: Hainich-Dün; blue: Swabian Alb). **A:** of the GBS analysis and **B:** of the mRNA-GBS analysis (n_m: not mown, m: mown)

## Discussion

The ability to link population genetic and functional genomic analysis in a rapid, cost-effective, and technically relatively simple manner would be of great importance for a better understanding of naturally occurring variability and for breeding studies. This would allow for the simultaneous screening of diversity while identifying expression patterns and specific candidate genes involved in the response to certain species-specific environmental interactions. Currently this is very time consuming and costly (Bhat et al. 2016). Therefore, the method presented here, mRNA-GBS, aims to fill the gap by offering a low-cost reduced complexity transcriptome analysis (mRNA-GBS).

This is the first approach, linking a complexity reduced mRNA analysis (mRNA-GBS) with an in depth RNA-Seq analysis (Herbert et al. 2021) and a GBS approach on natural occurring plant populations and across a broader geographic scale. We tested the mRNA-GBS approach on several individuals of red clover from eleven populations and three regions in Germany. We hereby evaluate whether the analysis of intraspecific variation within and between populations and transcriptome responses is possible simultaneously. The mRNA-GBS approach revealed population genetic patterns, but linkage with mRNA-Seq data was not possible. The drawbacks and needed optimization steps are discussed in the following.

### mRNA-GBS and comparison with RNA-Seq and GBS

Herbert et al. (2021) conducted an RNA-Seq analysis on one of the here also screened populations of red clover from Hainich-Dün (H) to compare the global transcriptional response to mowing under greenhouse conditions and in agricultural fields. They simulated mowing and compared the transcriptome response in mown and not mown *T. pratense* individuals, as in our analysis. Herbert et al. (2021) obtained a total number of short reads ranging from 44.7 to 58.1 million for each library, which on average is 10-times more per individual than in our study. Their sequencing approach comprised 608.041.012 raw reads for the analysis of only six different sites/treatments, of eight pooled samples while in our mRNA-GBS approach we investigated 13 plants on five to six fields in three regions in Germany. With this approach, they were able to identify 119 – 142 differentially expressed genes (DEGs, with a log2fold-change <u>>2</u>) that are up- or down-regulated when mown plants were compared with non-mown plants. The mRNA-GBS library was highly variable in terms of read depth per individual (80 bp on average), and pooling of samples did not allow us to correlate site-specific multifactorial influences of environmental responses in a statistically robust way. Only 50-86 % of the retrieved short sequences are located within the 100 bp region upstream of the poly(A) tail, and only 0.9 – 3-2 % are located within the last 25 bp, which hampered mRNA mapping and prevented the screening for differentially expressed genes (Table S1). SNP calling and expression studies were thus not possible.

However, also Herbert et al. (2021) discovered that plants grown in the field exhibited more and different stress responses than plants grown in greenhouses, leading them to conclude that field grown plants respond to multiple environmental stresses that are of site specific, abiotic, and biotic in origin. For example, they found some genes upregulated in mown plants being chitinase homologs suggesting that these plants are stressed by insects and/or fungi and that this stress may be more relevant to the plants than the loss of biomass due to mowing. With more than 65 different fungi and nematodes and more than 20 viruses, insects, and bacteria known to infect red clover (Duke 1981), our pilot study of mRNA-GBS across such a broad geographic and ecologically diverse range was too ambitious.

Our sequencing depth with an average of 1.1 million raw reads per sample for mRNA-GBS was too shallow to quantify gene expression differences. Hou et al (2013) proposed sequencing of 15-50 million reads to allow the detection of the majority of transcripts in human tissue (1C value between 2.9-3.1, Lander et al. 2001), so that a 15-fold higher read depth must be aimed for, which, however, does not meet our requirements that the method be inexpensive and easy to perform on multiple individuals. However, the high error rates resulting from the low sequencing depth are due to conceptual and methodological limitations of NGS sequencing, resulting in artifacts and a relatively high false positive rate of variants such as SNPs and InDels, not only affect the mRNA-GBS approach but also estimates of population genetic parameters (Dorant et al., 2019; Andrews et al., 2016; Cariou et al., 2016; Davey et al., 2011). This became apparent when we pooled the different individuals from mown and not mown populations from the mRNA-GBS analysis from each region, mapped them against a reference genome, and analyzed SNPs and compared them to the GBS analysis. The genetic diversity indices revealed significant inconsistencies between H_e_-GBS (ØH_e_ = 0.060) and H_e_-mRNA-GBS (ØH_e_ = 0.76) values. Our inconsistencies are based on the fact that different evolutionary mechanisms exert both neutral processes such as drift and immigration and adaptive processes such as selection, so that the different evolutionary origins of SNPs limit significance and may also overlap signals (Lamy et al., 2017; Vellend & Geber, 2005). Furthermore, Dorant et al. (2019) previously pointed out problems associated with GBS involving mutations at restriction sites that lead to allelic dropouts and PCR biases such that correct genetic diversity is not reflected and significant misinterpretation of commonly used statistics in population genetics studies leads to incorrect conclusions (Arnold et al., 2013, Cariou et al., 2016; Gautier et al., 2015). Several studies investigated the genetic diversity of red clover populations and germplasm collections, e.g., using RAPD (Campos-de-Quiroz & Ortega-Klose 2001; Ulloa et al. 2003), AFLP (Kölliker et al. 2003; Herrmann et al., 2005), and SSR (Gupta et al. 2017), and several of them found relatively high values for genetic diversity estimates similar to or slightly lower than those of our mRNA-GBS analysis. Pfeifer et al. (2018) compared GBS and AFLP data in an herbaceous perennial sedge species (*Carex gayana*) and found slightly higher estimates of genetic diversity with SNPs than with AFLP data, but also discovered some populations where this trend was reversed. SNP mutation rates are relatively low (10 × 10-8 to 10 × 10-9; Nachman & Crowell 2000; Pfeifer et al. 2018), lower than those of microsatellites (0.001 to 0.005; Pinto et al. 2013; Fischer et al. 2017), whereas AFLP mutation rates can exceed those of microsatellites (Kuchma et al. 2011).

STRUCTURE analyses revealed two genetic clusters for the GBS and pooled mRNA-GBS results, and the patterns were nearly region-specific in both analyses. In the mRNA-GBS, they were even treatment-specific (mown/not mown), which is weakly supported by PCA (Figs. 5) but no longer evident in the Neighbor Joining analysis. Deeper sequencing would potentially lead to the detection of mRNA sequences with lower copy number, resulting in stronger site-specific pattern recognition. GBS analysis revealed greater genetic similarity between individuals from S and H and a greater distance from A, with greater overlap in the mRNA-GBS data when all loci were considered; only at polymorphic loci did this pattern disappear. This is consistent with the results of other population genetic comparisons of plants studied in the Biodiversity Exploratories, e.g., *Veronica chamaedrys* (Kloss et al. 2011) in an AFLP study. While Kloss et al. (2011) found very little difference within and between populations, suggesting that the effects of genetic drift are counterbalanced by gene flow between populations, we found some differences. Both red clover and *V. chamaedrys* are commonly outcrossing perennials for which high gene flow is known to counteract the effects of genetic drift, either through high natural or human-induced dispersal of seeds and pollen or through large effective population sizes (Nybom 2004; Musche et al. 2008).

### mRNA-GBS and other marker assisted approaches

The advantage of mRNA-GBS is that it provides SNPs of transcripts from very specific biological processes at a specific time point and under the conditions prevailing there that characterize the phenotype, even if we mainly target the far 3’ end. In contrast, the GBS approach and similar molecular techniques used for NGS-based population genomic analyses (e.g., Hy-Rad, ddRAD-Seq, Pool-Seq, Hy-Rad, restriction site-associated DNA capture (Rapture), bulk and low-coverage NGS, and others, e.g., discussed in Dorant et al. 2019) provide SNPs from genomic regions and reflect only genotype, whereas phenotype is influenced by both its genotype and environment. RNA-Seq experiments targeting the phenotype can currently only be performed for a limited number of individuals and replicates due to the high cost of library preparation and deep sequencing, and assignment to a reference genome is required (Pallares et al. 2020). For marker assisted breeding as well as a better understanding of natural variability in populations the mRNA-GBS approach aim to identify specific traits through the use of direct and indirect molecular markers to replace standard comparative in-depth transcriptomics (Collard & Makill 2008).

Currently one approach is published, investigating gene expression in non-model plant populations with reduced complexities (Marx et al. 2020) and in comparison with RNA-Seq by using a TagSeq approach. Marx et al. (2020) performed RNA-Seq analysis on four non-model species at their natural populations. They then mapped TagSeq data from individuals at weekly intervals over three weeks and were able to align the short sequences with the reference transcriptome. However, they did not analyze these findings in an population genetic context. The TM3’seq approach (Pallares et al. 2020) also targets 3’ ends of transcripts while preserving sample identity at each step and enables simultaneous high-throughput processing of individual samples, but this approach has not been explored on plant samples, yet.

## Conclusion

In summary, we found that mRNA-GBS is a promising tool for population genetic analysis, but greater sequencing depth is required and fewer divergent populations need to be compared. The mRNA-GBS analysis described here resulted in too many divergent short sequence reads throughout the mRNA, making assignment difficult. It is recommended to focus more on generating mRNA regions upstream of the poly(a) tail. Experimental bias occurred in our analysis due to the use of NGS and GBS tools, which were pointed out previously. However, relative similarity and comparability of population genetic analysis is given, with mRNA-GBS data reflecting stronger signals of selection than neutral mutations compared with GBS data. Our approach has contributed to knowledge enhancement at a time when intensive research on genomic fingerprinting analyses and reduced RNA-Seq approaches is underway, particularly for non-model species

## Supporting information

Table S1 mRNA-GBS reads in correlation to mRNA position near the poly(A) region

## Acknowledgements

We thank the managers of the three Exploratories and all former managers, for their work in maintaining the plot and project infrastructure, Christiane Fischer and Jule Mangels for their support through the central office, Andreas Ostrowski and Michael Owonibi for managing the central database, and Markus Fischer, Eduard Linsenmair, Dominik Hessenmöller, Daniel Prati, Ingo Schöning, Francois Buscot, Ernst-Detlef Schulze, Wolfgang Weisser and the late Elisabeth Kalko for their role in setting up the Biodiversity Exploratories project. Fieldwork permits were issued by the responsible state environmental offices of Baden-Württemberg, Thüringen, and Brandenburg (according to § 72 331 BbgNatSchG). We are grateful to Volker Wissemann, Sabine Mutz, Annalena Kurzweil, Dr. Thomas Groß and Andreas Kolter for lab and administrative support. We are thankful to Andrea Weisert for carrying out all RNA extractions and cDNA synthesis steps.

