## Supplementary material for "GBS and a newly developed mRNA-GBS approach to link population genetic and transcriptome analyses reveal pattern differences between sites and treatments in red clover (*Trifolium pratense* L.)": Table S1 mRNA-GBS reads in correlation to mRNA position near the poly(A) region

| Sample | full length | % | 100 last bp close to poly(A) | % | 75 last bp close to poly(A) | % | 50 last bp close to poly(A) | % | 25 last bp close to poly(A) | % |
|---|---|---|---|---|---|---|---|---|---|---|
| A14-1 | 329586 | 67,0 | 13941 | 2,8 | 12337 | 2,5 | 10447 | 2,1 | 8156 | 1,7 |
| A14-2 | 268314 | 69,5 | 12040 | 3,1 | 10524 | 2,7 | 8895 | 2,3 | 6728 | 1,7 |
| A14-3 | 200873 | 60,3 | 10849 | 3,3 | 9393 | 2,8 | 8040 | 2,4 | 6291 | 1,9 |
| A14-4 | 202126 | 71,1 | 8571 | 3,0 | 7437 | 2,6 | 6202 | 2,2 | 4604 | 1,6 |
| A14-5 | 134723 | 67,1 | 7181 | 3,6 | 6131 | 3,1 | 5256 | 2,6 | 3757 | 1,9 |
| A14-6 | 253199 | 59,9 | 11395 | 2,7 | 10030 | 2,4 | 8389 | 2,0 | 6655 | 1,6 |
| A14-7 | 132196 | 74,0 | 5392 | 3,0 | 4582 | 2,6 | 3741 | 2,1 | 2752 | 1,5 |
| A15-1 | 194106 | 66,5 | 6031 | 2,1 | 5150 | 1,8 | 4245 | 1,5 | 2965 | 1,0 |
| A15-2 | 560596 | 83,0 | 39085 | 5,8 | 34384 | 5,1 | 28859 | 4,3 | 21419 | 3,2 |
| A15-3 | 288213 | 72,0 | 16916 | 4,2 | 14762 | 3,7 | 12556 | 3,1 | 9978 | 2,5 |
| A15-4 | 229961 | 50,8 | 19859 | 4,4 | 17627 | 3,9 | 15178 | 3,4 | 12266 | 2,7 |
| A15-5 | 258432 | 77,2 | 18250 | 5,4 | 15851 | 4,7 | 13212 | 3,9 | 9952 | 3,0 |
| A15-6 | 280579 | 74,5 | 9460 | 2,5 | 8081 | 2,1 | 6570 | 1,7 | 4952 | 1,3 |
| A15-7 | 327355 | 70,3 | 9071 | 1,9 | 7962 | 1,7 | 6374 | 1,4 | 4810 | 1,0 |
| A2-1 | 154629 | 63,7 | 6160 | 2,5 | 5408 | 2,2 | 4565 | 1,9 | 3438 | 1,4 |
| A2-2 | 137457 | 68,7 | 4782 | 2,4 | 4138 | 2,1 | 3440 | 1,7 | 2530 | 1,3 |
| A2-3 | 215731 | 67,4 | 9328 | 2,9 | 7966 | 2,5 | 6644 | 2,1 | 5033 | 1,6 |
| A24-1 | 212818 | 68,9 | 9845 | 3,2 | 8548 | 2,8 | 7032 | 2,3 | 5219 | 1,7 |
| A24-2 | 224742 | 63,8 | 9291 | 2,6 | 8182 | 2,3 | 7106 | 2,0 | 5636 | 1,6 |
| A24-3 | 202917 | 64,8 | 7582 | 2,4 | 6364 | 2,0 | 5179 | 1,7 | 3825 | 1,2 |
| A2-4 | 43589 | 60,9 | 3524 | 4,9 | 3060 | 4,3 | 2658 | 3,7 | 2108 | 2,9 |
| A24-4 | 56027 | 68,4 | 1557 | 1,9 | 1287 | 1,6 | 1064 | 1,3 | 810 | 1,0 |
| A24-5 | 213867 | 68,5 | 10551 | 3,4 | 9335 | 3,0 | 7895 | 2,5 | 5816 | 1,9 |
| A24-6 | 170648 | 56,3 | 9829 | 3,2 | 8462 | 2,8 | 7303 | 2,4 | 5751 | 1,9 |
| A24-7 | 183259 | 71,4 | 8533 | 3,3 | 7351 | 2,9 | 6066 | 2,4 | 4384 | 1,7 |
| A2-5 | 72955 | 68,3 | 2004 | 1,9 | 1720 | 1,6 | 1420 | 1,3 | 995 | 0,9 |
| A2-6 | 116445 | 69,1 | 3465 | 2,1 | 2933 | 1,7 | 2396 | 1,4 | 1661 | 1,0 |
| A2-7 | 158311 | 66,9 | 5872 | 2,5 | 5132 | 2,2 | 4252 | 1,8 | 3146 | 1,3 |
| A31-1 | 161172 | 70,7 | 6007 | 2,6 | 5140 | 2,3 | 4188 | 1,8 | 3060 | 1,3 |
| A31-2 | 106411 | 74,7 | 4613 | 3,2 | 3893 | 2,7 | 3161 | 2,2 | 2310 | 1,6 |
| A31-3 | 293516 | 72,4 | 12826 | 3,2 | 10861 | 2,7 | 8797 | 2,2 | 6263 | 1,5 |
| A31-4 | 234012 | 74,5 | 10809 | 3,4 | 9184 | 2,9 | 7597 | 2,4 | 5564 | 1,8 |
| A31-5 | 158155 | 75,7 | 6988 | 3,3 | 5915 | 2,8 | 4880 | 2,3 | 3693 | 1,8 |
| A31-6 | 45910 | 80,6 | 2597 | 4,6 | 2209 | 3,9 | 1785 | 3,1 | 1298 | 2,3 |
| A31-7 | 80351 | 72,9 | 5112 | 4,6 | 4201 | 3,8 | 3527 | 3,2 | 2719 | 2,5 |
| A9-1 | 254261 | 80,9 | 8649 | 2,8 | 7241 | 2,3 | 5496 | 1,7 | 4021 | 1,3 |
| A9-2 | 119989 | 77,8 | 4407 | 2,9 | 3594 | 2,3 | 2636 | 1,7 | 1875 | 1,2 |
| A9-3 | 351520 | 80,0 | 11978 | 2,7 | 10023 | 2,3 | 7777 | 1,8 | 5634 | 1,3 |
| A9-4 | 278663 | 72,9 | 11623 | 3,0 | 10004 | 2,6 | 8059 | 2,1 | 5936 | 1,6 |
| A9-5 | 274145 | 67,8 | 13263 | 3,3 | 11332 | 2,8 | 9313 | 2,3 | 7207 | 1,8 |
| A9-6 | 83444 | 87,9 | 5082 | 5,4 | 4238 | 4,5 | 3385 | 3,6 | 2514 | 2,6 |
| A9-7 | 357633 | 83,8 | 15681 | 3,7 | 13283 | 3,1 | 10229 | 2,4 | 7705 | 1,8 |
| H13-1 | 249411 | 63,8 | 10696 | 2,7 | 9629 | 2,5 | 7912 | 2,0 | 5641 | 1,4 |
| H13-2 | 157436 | 72,9 | 6941 | 3,2 | 6045 | 2,8 | 4966 | 2,3 | 3571 | 1,7 |
| H13-3 | 271452 | 67,8 | 13311 | 3,3 | 11785 | 2,9 | 9778 | 2,4 | 7385 | 1,8 |
| H13-4 | 256185 | 71,7 | 9291 | 2,6 | 8179 | 2,3 | 6678 | 1,9 | 4809 | 1,3 |

| | | | | | | | | | | |
|---|---|---|---|---|---|---|---|---|---|---|
| H13-5 | 262617 | 70,9 | 14121 | 3,8 | 12994 | 3,5 | 11598 | 3,1 | 10215 | 2,8 |
| H13-6 | 238938 | 71,0 | 10228 | 3,0 | 9003 | 2,7 | 7410 | 2,2 | 5223 | 1,6 |
| H13-7 | 195323 | 67,7 | 7918 | 2,7 | 7033 | 2,4 | 5683 | 2,0 | 4093 | 1,4 |
| H15-1 | 219733 | 68,6 | 5352 | 1,7 | 4505 | 1,4 | 3754 | 1,2 | 3121 | 1,0 |
| H15-2 | 32130 | 62,9 | 1943 | 3,8 | 1663 | 3,3 | 1335 | 2,6 | 1059 | 2,1 |
| H15-3 | 13157 | 66,9 | 431 | 2,2 | 358 | 1,8 | 287 | 1,5 | 220 | 1,1 |
| H15-4 | 118760 | 58,4 | 8128 | 4,0 | 7182 | 3,5 | 6080 | 3,0 | 4718 | 2,3 |
| H15-5 | 246482 | 76,4 | 10128 | 3,1 | 8850 | 2,7 | 7014 | 2,2 | 5069 | 1,6 |
| H15-6 | 79181 | 71,3 | 3558 | 3,2 | 3087 | 2,8 | 2469 | 2,2 | 1868 | 1,7 |
| H15-7 | 117768 | 73,0 | 6409 | 4,0 | 5590 | 3,5 | 4465 | 2,8 | 3337 | 2,1 |
| H17-1 | 1541831 | 72,0 | 63016 | 2,9 | 54521 | 2,5 | 45304 | 2,1 | 30782 | 1,4 |
| H17-2 | 20977 | 55,2 | 1781 | 4,7 | 1578 | 4,2 | 1351 | 3,6 | 1094 | 2,9 |
| H17-3 | 42820 | 66,5 | 3063 | 4,8 | 2626 | 4,1 | 2181 | 3,4 | 1726 | 2,7 |
| H17-4 | 299819 | 74,7 | 16795 | 4,2 | 14051 | 3,5 | 11735 | 2,9 | 8311 | 2,1 |
| H17-5 | 386320 | 69,7 | 23454 | 4,2 | 19649 | 3,5 | 16729 | 3,0 | 11419 | 2,1 |
| H17-6 | 198945 | 75,0 | 12089 | 4,6 | 10517 | 4,0 | 8908 | 3,4 | 5683 | 2,1 |
| H17-7 | 16411 | 58,6 | 943 | 3,4 | 808 | 2,9 | 697 | 2,5 | 526 | 1,9 |
| H50-1 | 1213721 | 73,9 | 59511 | 3,6 | 51716 | 3,1 | 43781 | 2,7 | 27321 | 1,7 |
| H50-2 | 726680 | 75,3 | 34191 | 3,5 | 29306 | 3,0 | 24197 | 2,5 | 16811 | 1,7 |
| H50-3 | 1906741 | 74,0 | 76988 | 3,0 | 68431 | 2,7 | 56970 | 2,2 | 35177 | 1,4 |
| H50-4 | 1357972 | 74,4 | 83424 | 4,6 | 71868 | 3,9 | 60832 | 3,3 | 41178 | 2,3 |
| H50-5 | 101585 | 74,4 | 3382 | 2,5 | 2590 | 1,9 | 2060 | 1,5 | 1531 | 1,1 |
| H50-6 | 927144 | 73,7 | 51583 | 4,1 | 45726 | 3,6 | 38467 | 3,1 | 24737 | 2,0 |
| H50-7 | 1050652 | 73,8 | 56381 | 4,0 | 49641 | 3,5 | 41551 | 2,9 | 26998 | 1,9 |
| H50g-1 | 1035201 | 72,6 | 54330 | 3,8 | 47909 | 3,4 | 40545 | 2,8 | 25113 | 1,8 |
| H50g-2 | 147714 | 68,3 | 6932 | 3,2 | 6148 | 2,8 | 5214 | 2,4 | 3091 | 1,4 |
| H50g-3 | 493846 | 69,3 | 18407 | 2,6 | 15962 | 2,2 | 13394 | 1,9 | 8539 | 1,2 |
| H50g-4 | 348258 | 75,3 | 14687 | 3,2 | 12375 | 2,7 | 10111 | 2,2 | 7101 | 1,5 |
| H50g-5 | 580340 | 67,7 | 29554 | 3,4 | 26085 | 3,0 | 22078 | 2,6 | 13750 | 1,6 |
| H50g-6 | 115171 | 74,2 | 3280 | 2,1 | 2538 | 1,6 | 2002 | 1,3 | 1423 | 0,9 |
| H50g-7 | 822043 | 74,8 | 34065 | 3,1 | 29321 | 2,7 | 24018 | 2,2 | 16948 | 1,5 |
| H8-1 | 194104 | 75,2 | 12423 | 4,8 | 10816 | 4,2 | 8481 | 3,3 | 6471 | 2,5 |
| H8-2 | 152486 | 63,8 | 6701 | 2,8 | 5789 | 2,4 | 4689 | 2,0 | 3553 | 1,5 |
| H8-3 | 113807 | 64,5 | 5759 | 3,3 | 4938 | 2,8 | 4107 | 2,3 | 3232 | 1,8 |
| H8-4 | 203464 | 79,6 | 9601 | 3,8 | 8204 | 3,2 | 6443 | 2,5 | 4877 | 1,9 |
| H8-5 | 216544 | 61,5 | 11705 | 3,3 | 10342 | 2,9 | 8704 | 2,5 | 6842 | 1,9 |
| H8-6 | 277115 | 74,2 | 11489 | 3,1 | 9989 | 2,7 | 7958 | 2,1 | 5830 | 1,6 |
| H8-7 | 187800 | 73,8 | 5085 | 2,0 | 4324 | 1,7 | 3532 | 1,4 | 2602 | 1,0 |
| S30-1 | 381956 | 65,9 | 16674 | 2,9 | 14838 | 2,6 | 12461 | 2,1 | 7302 | 1,3 |
| S30-2 | 238303 | 67,8 | 7426 | 2,1 | 6358 | 1,8 | 5433 | 1,5 | 4366 | 1,2 |
| S30-3 | 361037 | 67,9 | 13810 | 2,6 | 12061 | 2,3 | 10267 | 1,9 | 6681 | 1,3 |
| S30-4 | 697582 | 70,0 | 33462 | 3,4 | 29428 | 3,0 | 24783 | 2,5 | 16674 | 1,7 |
| S30-5 | 592301 | 68,7 | 23331 | 2,7 | 20438 | 2,4 | 17177 | 2,0 | 11779 | 1,4 |
| S30-6 | 724627 | 67,2 | 34124 | 3,2 | 30258 | 2,8 | 25687 | 2,4 | 17116 | 1,6 |
| S30-7 | 1136351 | 66,5 | 44581 | 2,6 | 39542 | 2,3 | 33467 | 2,0 | 21258 | 1,2 |
| S31-1 | 419624 | 65,6 | 20491 | 3,2 | 18183 | 2,8 | 15542 | 2,4 | 9891 | 1,5 |
| S31-2 | 659047 | 73,3 | 36669 | 4,1 | 31986 | 3,6 | 26857 | 3,0 | 18688 | 2,1 |
| S31-3 | 306787 | 67,1 | 15218 | 3,3 | 13451 | 2,9 | 11500 | 2,5 | 7367 | 1,6 |
| S31-4 | 692772 | 65,7 | 34912 | 3,3 | 31327 | 3,0 | 26853 | 2,5 | 16894 | 1,6 |
| S31-5 | 362795 | 71,5 | 10946 | 2,2 | 8753 | 1,7 | 6954 | 1,4 | 4892 | 1,0 |

| | | | | | | | | | | |
|---|---|---|---|---|---|---|---|---|---|---|
| S31-6 | 1780605 | 71,3 | 93612 | 3,7 | 82466 | 3,3 | 69769 | 2,8 | 48885 | 2,0 |
| S31-7 | 1372862 | 62,1 | 53632 | 2,4 | 47367 | 2,1 | 39997 | 1,8 | 27310 | 1,2 |
| S32-1 | 230715 | 67,7 | 8830 | 2,6 | 7692 | 2,3 | 6504 | 1,9 | 3713 | 1,1 |
| S32-2 | 517720 | 74,2 | 19186 | 2,8 | 16100 | 2,3 | 13152 | 1,9 | 9055 | 1,3 |
| S32-3 | 315614 | 68,3 | 9512 | 2,1 | 7944 | 1,7 | 6664 | 1,4 | 5046 | 1,1 |
| S32-4 | 651637 | 64,9 | 31192 | 3,1 | 27513 | 2,7 | 23185 | 2,3 | 15404 | 1,5 |
| S32-5 | 159330 | 71,5 | 6824 | 3,1 | 5914 | 2,7 | 4935 | 2,2 | 3577 | 1,6 |
| S32-6 | 325683 | 65,7 | 16060 | 3,2 | 14169 | 2,9 | 12008 | 2,4 | 7977 | 1,6 |
| S32-7 | 632191 | 71,4 | 20705 | 2,3 | 17462 | 2,0 | 14352 | 1,6 | 10413 | 1,2 |
| SHG-1 | 703688 | 58,3 | 52116 | 4,3 | 45937 | 3,8 | 39375 | 3,3 | 30947 | 2,6 |
| SHG-2 | 922139 | 69,5 | 55872 | 4,2 | 48602 | 3,7 | 40629 | 3,1 | 26825 | 2,0 |
| SHG-3 | 502073 | 81,0 | 36254 | 5,9 | 31348 | 5,1 | 26492 | 4,3 | 19745 | 3,2 |
| SHG-4 | 390460 | 57,3 | 29037 | 4,3 | 25370 | 3,7 | 21812 | 3,2 | 17032 | 2,5 |
| SHG-5 | 211484 | 75,6 | 10643 | 3,8 | 9224 | 3,3 | 7651 | 2,7 | 4938 | 1,8 |
| SHG-6 | 279430 | 75,7 | 15213 | 4,1 | 13162 | 3,6 | 11054 | 3,0 | 7397 | 2,0 |
| SHG-7 | 15878 | 64,7 | 632 | 2,6 | 512 | 2,1 | 386 | 1,6 | 276 | 1,1 |
| Sz1-1 | 1586227 | 63,0 | 116660 | 4,6 | 103216 | 4,1 | 87890 | 3,5 | 66148 | 2,6 |
| Sz1-2 | 279941 | 76,5 | 10039 | 2,7 | 8281 | 2,3 | 6384 | 1,7 | 4496 | 1,2 |
| Sz1-3 | 875633 | 72,6 | 42392 | 3,5 | 37114 | 3,1 | 31763 | 2,6 | 20310 | 1,7 |
| Sz1-4 | 927901 | 74,7 | 45844 | 3,7 | 40051 | 3,2 | 33806 | 2,7 | 21434 | 1,7 |
| Sz1-5 | 791205 | 73,5 | 36557 | 3,4 | 31947 | 3,0 | 26722 | 2,5 | 16648 | 1,5 |
| Sz1-6 | 626613 | 76,8 | 34658 | 4,2 | 30427 | 3,7 | 25474 | 3,1 | 17708 | 2,2 |
| Sz1-7 | 39427 | 62,6 | 1836 | 2,9 | 1527 | 2,4 | 1230 | 2,0 | 948 | 1,5 |
| Sz2-1 | 14158 | 68,0 | 441 | 2,1 | 351 | 1,7 | 267 | 1,3 | 179 | 0,9 |
| Sz2-2 | 277851 | 76,4 | 16958 | 4,7 | 14030 | 3,9 | 11741 | 3,2 | 8223 | 2,3 |
| Sz2-3 | 156946 | 77,1 | 9173 | 4,5 | 7936 | 3,9 | 6301 | 3,1 | 4059 | 2,0 |
| Sz2-4 | 829546 | 74,6 | 42250 | 3,8 | 36707 | 3,3 | 30347 | 2,7 | 20484 | 1,8 |
| Sz2-5 | 266728 | 71,8 | 8826 | 2,4 | 7185 | 1,9 | 5870 | 1,6 | 4278 | 1,2 |
| Sz2-6 | 60793 | 78,5 | 2015 | 2,6 | 1592 | 2,1 | 1215 | 1,6 | 873 | 1,1 |
| Sz2-7 | 35494 | 74,2 | 2119 | 4,4 | 1759 | 3,7 | 1370 | 2,9 | 1038 | 2,2 |
| total | 48114074 | | 2339805 | | 2038454 | | 1706608 | | 1186705 | |
| on average | 381858 | 70,23 | 18570 | 3,31 | 16178 | 2,86 | 13545 | 2,37 | 9418 | 1,71 |
| min | 13157 | 50,8 | 431 | 1,7 | 351 | 1,4 | 267 | 1,2 | 179 | 0,9 |
| max | 1906741 | 87,9 | 116660 | 5,9 | 103216 | 5,1 | 87890 | 4,3 | 66148 | 3,2 |
